# Glial activation in prion diseases is strictly nonautonomous and requires neuronal PrP^Sc^

**DOI:** 10.1101/2021.01.03.425136

**Authors:** Asvin KK Lakkaraju, Silvia Sorce, Assunta Senatore, Mario Nuvolone, Jingjing Guo, Petra Schwarz, Rita Moos, Pawel Pelczar, Adriano Aguzzi

## Abstract

Although prion infections cause cognitive impairment and neuronal death, transcriptional and translational profiling shows progressive derangement within glia but surprisingly little changes within neurons. Here we expressed PrP^C^ selectively in neurons, astrocytes or oligodendrocytes of mice. After prion infection, both astrocyte and neuron-restricted PrP^C^ expression led to copious brain accumulation of PrP^Sc^. As expected, neuron-restricted expression was associated with typical prion disease. However, mice with astrocyte-restricted PrP^C^ expression experienced a normal life span, did not develop clinical disease, and did not show astro- or microgliosis. Besides confirming that PrP^Sc^ is innocuous to PrP^C^-deficient neurons, these results show that astrocyte-born PrP^Sc^ does not activate the extreme neuroinflammation that accompanies the onset of prion disease and precedes any molecular changes of neurons. This points to a nonautonomous mechanism by which prion-infected neurons instruct astrocytes and microglia to acquire a specific cellular state that, in turn, drives neural dysfunction.

## Introduction

Prion diseases are characterized by a long, largely asymptomatic incubation period. Once clinical signs and symptoms arise, the disease typically progresses very rapidly. Prion-infected brains contain PrP^Sc^, an aggregated and misfolded isoform of the cellular prion protein (PrP^C^) (Aguzzi, 2009). PrP^Sc^ seeds the nucleation of prions by recruiting PrP^C^; accordingly, ablation of PrP^C^ abrogates prion propagation (Bueler et al., 1993) and toxicity (Brandner et al., 1996).

The clinical manifestation of prion diseases, both in humans and in animal models, consist of progressive neurological signs including deterioration of cortical functions. The anatomical correlates of the disease are spongiosis (a peculiar form of neuronal vacuolation), activation and proliferation of microglia, and astrogliosis. The cortex of patients with terminal prion diseases can show an almost total depletion of neurons (Aguzzi et al., 2013; Budka, 2003), which suggests that neurons may be the primary target of the disease. But what is the connection between prion replication and neuronal demise? In a neurografting paradigm, *Prnp*-ablated neurons survive long-term exposure to prions (Brandner et al., 1996), implying that resident PrP^C^ is necessary for the development of damage. Also, quenching neuronal PrP^C^ expression was found to delay prion disease (Mallucci et al., 2003), adding to the evidence that neuronal PrP^C^ is required for neurotoxicity.

However, recent molecular studies are painting a starkly different picture. Transcriptomic analysis performed in prion-infected mice over the course of disease have revealed dramatic aberrations of glia-enriched genes coinciding with the onset of clinical signs, whereas neuronal changes were less pronounced and were only detected at the terminal stage of the disease (Sorce et al., 2020). Similarly, a quantitative analysis of mRNA translation during the course of prion diseases has found that almost all changes during the progression of prion disease occur in non-neuronal cells, except very late in disease (Scheckel et al., 2020). These findings suggest that it is the glia which experiences initial dysfunction, whereas the neuronal demise is a consequence thereof.

Here we have tested the above hypothesis by systematically investigating the impact of cell-type specific PrP^C^ in disease. We generated transgenic mice line expressing a conditionally expressed PrP transgene, and mated them with mice expressing the Cre recombinase driven by cell-type specific promoters. The resulting mice expressed PrP^C^ in a cell-type specific manner. We found that neuron-restricted PrP^C^ sufficed to rescue the demyelination defects observed in the PrP^C^ ablated mice and to induce neurodegeneration upon prion infection. Conversely, mice with astrocyte-restricted PrP^C^ expression did not experience clinical signs of scrapie after prion infection. However, we found that these mice had conspicuous PrP^Sc^ accumulation.

## Results

### Generation of mice with cell-type restricted PrP^C^ expression

We developed a versatile transgenic mouse model that directs expression to specific cell types in a robust and controllable manner. We used a murine *Prnp* cDNA under the transcriptional control of the cytomegalovirus/chicken beta actin/rabbit beta-globin gene (CAG) promoter and a loxP-stop-loxP (LSL) cassette flanking the chloramphenicol acetyl transferase (CAT) gene (Araki et al., 1995). When crossed to mice expressing the Cre recombinase tissue-specifically, the CAT gene and the transcriptional stop cassette flanked by LoxP sites are excised, permitting activation of PrP^C^ expression (Figure 1A).

**Figure 1:**
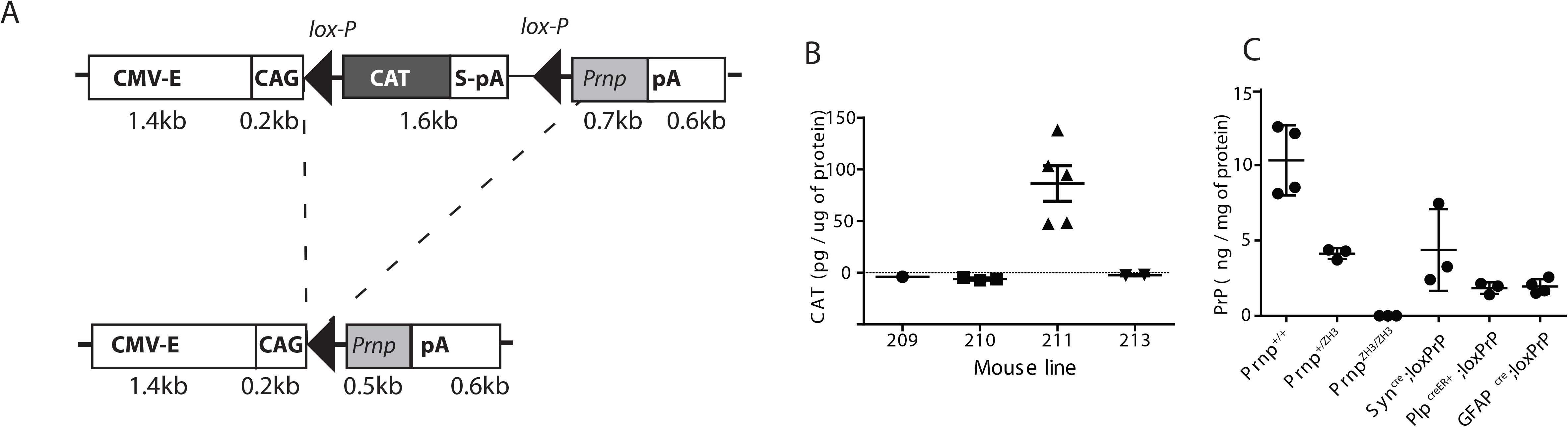
Characterization of mice expressing PrP in defined brain compartments. A: Schematic of the CAGCAT-Prnp transgene: The transgene is driven by the CMV enhancer followed by chicken beta actin (CAG)promoter. The chloramphenicol acetyl transferase (CAT) expression cassette contains an SV40 polyadenylation signal (S-pA) and is flanked by lox-P sites, followed by the Prnp coding sequence and a rabbit β-globin polyadenylation signal (pA). Transgenic Prnp is expressed only upon excision of the CAT stop sequence by the Cre recombinase. B: Transgenic CAT expression in brain homogenates, assessed by ELISA. Assays here and in panel C were performed as triplicates; plots represent mean ± s.e.m. Line 211 displaying the highest CAT expression was selected for crossing with PrnpZH3/ZH3 mice. C: The resulting PrnpZH3/ZH3;CAG-CAT-PrP mice were crossed with Cre driver lines. Brain PrPC expression was compared to Prnp+/+, Prnp+/ZH and PrnpZH3/ZH3 mice by ELISA. Crosses to SynCre, GFAPCre and PlpcreER+ yielded selective PrPC expression in subset of neurons, astrocytes and oligodendrocytes, respectively, at approximately one third of wild-type mouse brains.

Pronuclear injection of the CAG-CAT-PrP construct into one-cell stage fertilized embryos from mice ablated of cellular PrP^C^ (*Prnp*^ZH1/ZH1^) resulted in the generation of seven transgenic lines (Figure S1) of which four lines successfully transmitted the transgene to F1 generation. We next monitored the expression of CAT in brain lysates from offspring of all four transgenic lines using quantitative enzyme-linked immunosorbent assay (ELISA). Only line 211 showed sustained expression of CAT (Figure 1B). Line 211 was further bred to co-isogenic C57BL/6 *Prnp*-ablated mice (Prnp^ZH3/ZH3^) for 9 successive generations.

To evaluate the robustness of CAG-CAT system in generating cell type specific PrP^C^ expressors, we crossed Line 211 with mice expressing Cre under the control of the synapsin-1, glial fibrillary acidic protein (GFAP) and proteolipid (Plp) promoters in order to direct PrP^C^ expression selectively to a subset of neurons (Syn^Cre^;loxPrP), astrocytes (GFAP^Cre^;loxPrP) and oligodendrocytes (Plp^Cre^;loxPrP) respectively. Mice were tested by ear-biopsy PCR for the CAG-CAT-PrP and the appropriate Cre transgene, and brain lysates from double-positive mice were subjected to ELISA. Syn^Cre^;loxPrP mice showed a higher expression of PrP^C^ than GFAP^Cre^;loxPrP and Plp^Cre^;loxPrP mice in their brains. As expected, the overall protein levels of PrP^C^ were lower in brain lysates of mice expressing cell-type specific PrP^C^ than wild-type mice or hemizygous *Prnp*^ZH3/+^ mice (Figure 1C).

To assess the cell-type restricted expression of PrP^C^, we performed immunofluorescence on the cerebellar brain sections. Mice lacking PrP^C^ (*Prnp*^ZH3/ZH3^) and wild-type mice were used as negative and positive controls. In cerebellar sections of Syn^Cre^;loxPrP mice, PrP^C^ expression was exclusively observed in Purkinje cells. Unlike Syn^Cre^;loxPrP mice, co-staining with MAP-2, which labels mature neuronal population, revealed that PrP^C^ expression in wild-type mice showed a diffuse neuronal staining. *Prnp*^ZH3/ZH3^ mice did not show any staining of PrP^C^ (Figure 2A). While several glia drivers were shown to also label neuronal subpopulations (Gregorian et al., 2009), the GFAP-Cre driver used here was shown to exclusively label astrocytes (Gregorian et al., 2009). Co-staining with GFAP in cerebellar sections of GFAP^Cre^;loxPrP mice revealed astrocytic localization of PrP^C^. Finally, no expression of PrP^C^ was seen in the absence of Cre (Figure 2B). Plp driven Cre-recombinase expression was found to be restricted predominantly to mature oligodendrocyte population, however recent studies have contradicted this (Michalski et al., 2011). Co-staining of cerebellar sections with myelin associated glycoprotein (MAG), which labels the mature oligodendrocytes, from mice expressing Plp^Cre^;loxPrP, revealed that PrP^C^ was exclusively localized to the oligodendrocytes and more specifically restricted to the white matter (Figure 2C).

**Figure 2:**
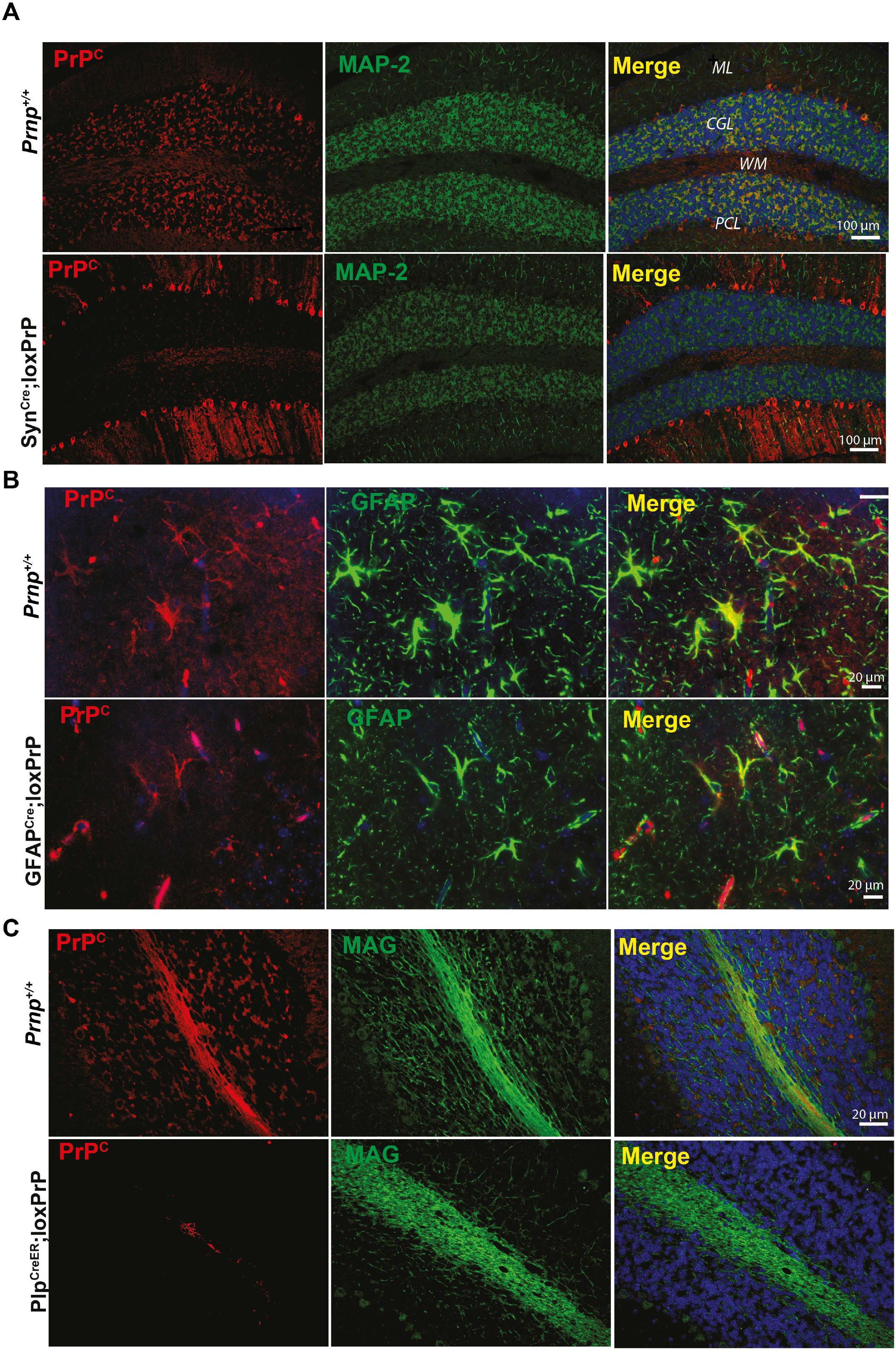
PrP^C^ expression patterns in CAG-CAT-PrP mice. **A-B:** Immunofluorescent staining of PrP^C^ (red) comparing cerebellar brain sections of wild-type Prnp^+/+^ mice to Syn^Cre^;loxPrP, GFAP^Cre^;loxPrP and Plp^creER^;loxPrP mice. Antibody to MAP2 (microtubule-associated protein 2), GFAP (glial fibrillary acidic protein) and MAG (myelin-associated glycoprotein) were used as neuronal, astrocytic and oligodendrocytic markers respectively (in green). Cell nuclei were stained with DAPI (in blue). In Prnp^+/+^ mice, PrP^C^ is detected in the cerebellar granule cell layer (CGL), Purkinje cells (PCL) and molecular layer (ML). In Syn^Cre^;loxPrP^+^ mice (A), PrP^C^ is only detected in Purkinje cells where the promoter controlling *Cre* is active. In GFAP^Cre^;loxPrP mice, PrP^C^ is exclusively detected in astrocytes where Cre is active. In Plp^creER^;loxPrP mice, PrP^C^ is only detected in the oligodendrocytes in the white matter colocalizing with MAG.

### Role of neuronal and astrocytic PrP^C^ in the manifestation of prion disease

We sought to investigate the kinetics and the manifestation of prion disease in the newly-generated mice expressing PrP^C^ in a subset of neurons (Zhu et al., 2001). Five Syn^Cre^;loxPrP mice were intracerebrally inoculated with prions (Rocky Mountain Laboratory strain, passage #6, henceforth termed RML6). For control we used wild-type C57BL6/J mice, mice transgenic for Syn^Cre^ but not for loxPrP, and vice versa. Syn^Cre^;loxPrP mice developed clinical scrapie, albeit with significantly longer incubation times than wild-type mice (435 vs 196 days after inoculation (dpi), respectively, *p*=0.002) (Figure 3A). Hence neuronal PrP^C^ expression suffices to confer susceptibility to prion disease. We next assessed proteinase K (PK) resistant prion protein (termed PrP^Sc^) in brain lysates of terminally scrapie-sick wild-type and Syn^Cre^;loxPrP mice, as well as age-matched loxPrP, Syn^Cre^ or *Prnp*^ZH3/ZH3^ mice. Western blotting of PK-treated lysates (25 μg/ml, 37°C, 1h) revealed PrP^Sc^ only in wild-type and Syn^Cre^;loxPrP lysates (Figure 3B). Next, five GFAP^Cre^;loxPrP transgenic mice were inoculated intracerebrally with RML6 prions. For control we used wild-type mice, GFAP^Cre^ mice, loxPrP mice, and *Prnp*^ZH3/ZH3^ mice. Over a period of 544 dpi, neither GFAP^Cre^;loxPrP mice nor *Prnp*^ZH3/ZH3^ mice developed any signs of scrapie, whereas *Prnp*^+/+^ mice (with or without Cre transgenes) developed terminal scrapie at 167 ± 5 dpi (Figure 3C). Western blots showed similar amounts of PrP^Sc^ in terminally scrapie-sick wild-type mice and in 2-year-old prion-infected GFAP^Cre^;loxPrP mice (544dpi) (Figure 3D).

**Figure 3:**
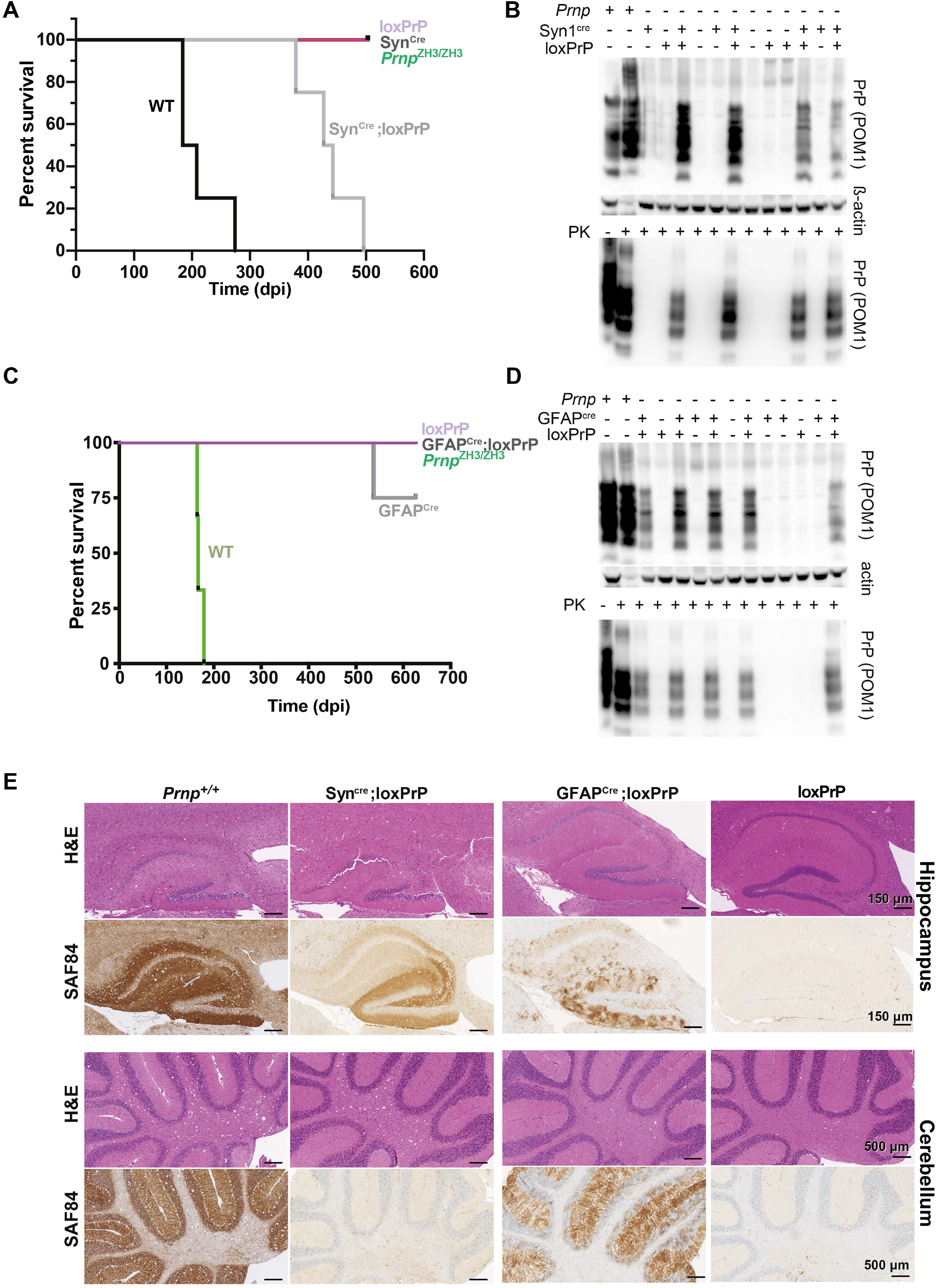
Neuron-restricted, but not astrocyte-restricted, PrP^C^ expression confers susceptibility to prion disease. **A:** Survival of Syn^Cre^;loxPrP, loxPrP, Syn^Cre^, Prnp^ZH3/ZH3^ and wild-type mice inoculated intracerebrally with RML6 prions. The median incubations for Syn^Cre^-;loxPrP^+^ (n = 5) and wild-type mice (n = 4) were 435 and 195 days post inoculation (dpi), respectively. Survival curves were compared by a log-rank (Mantel-Cox) test. **B:** Representative Western blot for total PrP (upper panel) and PK-resistant PrPSc (lower panel) in brains of prion-infected, terminally-sick mice. Syn^Cre^;loxPrP^+^ and wild-type brains contained PrP^Sc^. Detection: anti-PrP antibody POM1. β-actin: loading control. Lane #2 (upper panel) was intentionally underloaded in order to avoid overexposure. **C:** Survival curves of GFAP1^Cre^;loxPrP^+^, loxPrP, Prnp^ZH3/ZH3^ and wild-type mice inoculated with RML6 prions intracerebrally. GFAP^Cre^;loxPrP^+^ (n = 5) did not develop clinical signs of prion disease. Wildtype mice (n = 4): 167 ± 5 dpi. D: Western blot of total PrP and PK-digested PrP^Sc^ in brain homogenates of prion-infected GFAP^Cre^;loxPrP^+^ (626 dpi) and control mice. GFAP^Cre^;loxPrP^+^ harbor copious PrP^Sc^. E: Hippocampal and cerebellar histology of prion-infected Syn^Cre^;loxPrP^+^, GFAP^Cre^;loxPrP, loxPrP^-^ and Prnp^+/+^ mice. Slices were stained with haematoxylin and eosin (H&E) and anti-PrP antibody SAF84. Both astrocyte- and neuron-restricted PrP transgenic mice accumulated PrP^Sc^, but their deposition patterns differed profoundly.

Histological analysis of hippocampal and cerebellar brain sections revealed vacuolation (spongiosis) in Syn^Cre^;loxPrP mice, albeit less pronounced than in wild-type mice whereas, GFAP^Cre^;loxPrP did not show any vacuolation (Figure 3E). Immunohistochemistry with anti-PrP antibody SAF84 (Demart et al., 1999) in hippocampal sections of Syn^Cre^;loxPrP mice revealed the presence of PrP deposits, albeit at lower amounts than in wild-type mice, while mice lacking PrP or Cre-recombinase did not show any staining (Figure 3E). Syn^Cre^;loxPrP mice, which expressed PrP^C^ predominantly in Purkinje cells, exhibited showed fewer PrP deposits that wild-type mice (Figure 3E). SAF84 immunostaining in prion inoculated GFAP^Cre^;loxPrP mice however revealed PrP deposits in both cerebellum and hippocampus (Figure 3E). Taken together these results suggest that mice with neuron-restricted PrP^C^ develop clinical and histological features of prion disease whereas PrP^C^ expression in astrocytes does not restore prion-induced neurodegeneration, yet it supports PrP^Sc^ accumulation.

### Astrocyte restricted PrP^C^ does not induce neurodegeneration in COCS

We next generated cerebellar organotypic cultured slices (COCS) from 8-day old Syn^Cre^;loxPrP mice and inoculated them with RML6 prions or with non-infectious brain homogenate (NBH). At 56 dpi, COCS were immunostained with NeuN (staining cerebellar granule cells) and calbindin (staining Purkinje cells) (Weyer and Schilling, 2003).

Neuronal loss was measured by morphometric assessment of the area of cerebellar granule layer (CGL) immunoreactive to the antibodies against NeuN and calbindin. COCS generated from Syn^Cre^;loxPrP mice showed a significant decrease in calbindin staining, confirming neurodegeneration of PrP expressing Purkinje cells in the Syn^Cre^;loxPrP mice. Surprisingly, there was also a significant loss of afferent NeuN^+^ cerebellar granule neurons, possibly due to secondary effects arising from Purkinje cell death (Doughty et al., 2000). In contrast, control COCS from loxPrP mice did now show any neurodegeneration (Figure 4A).

**Figure 4:**
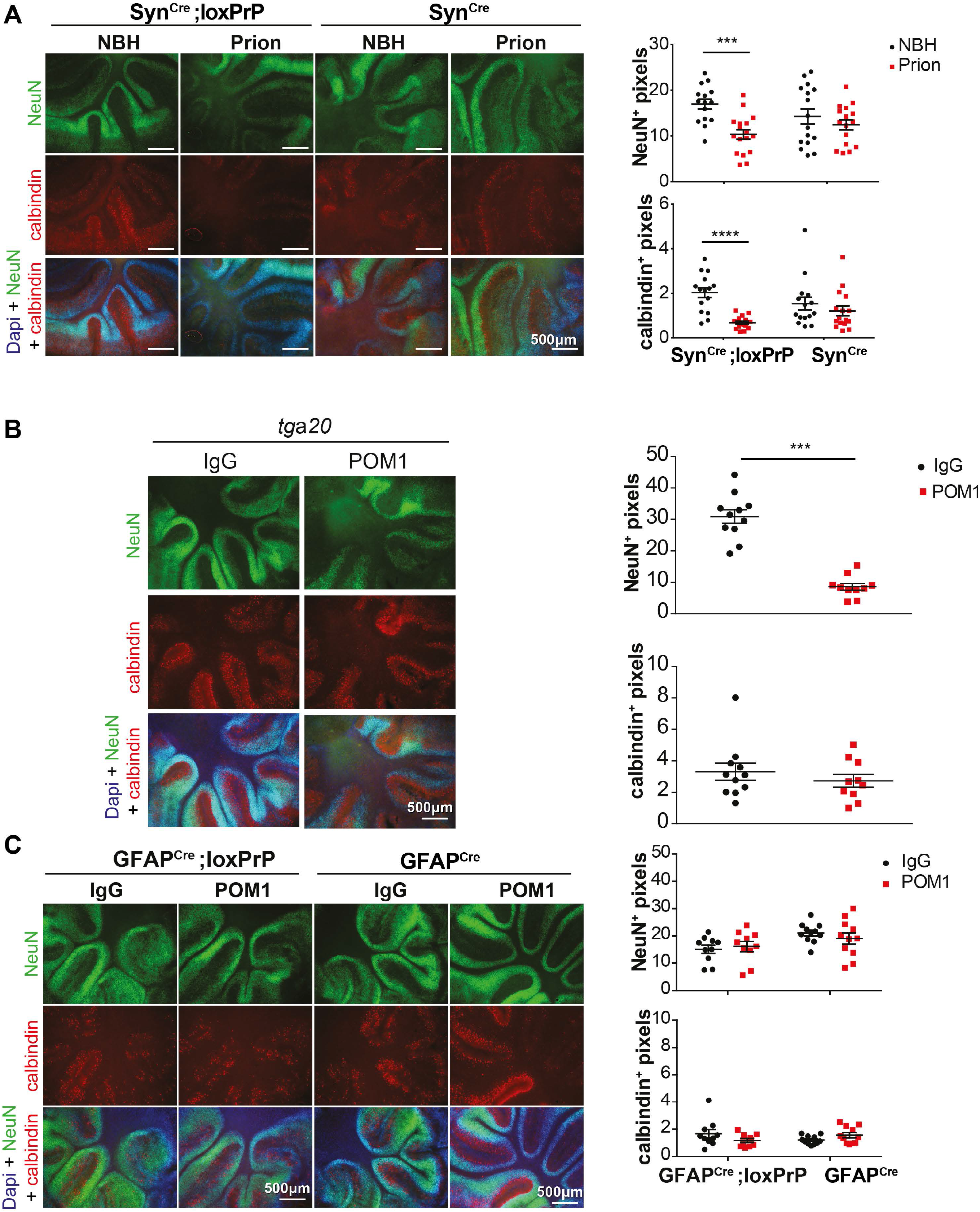
COCS from astrocyte-restricted PrP^C^ mice are resistant to neurodegeneration induced by prion mimetics. A: Neurodegeneration in prion-infected cerebellar organotypic brain slices (COCS) of Syn^Cre^;loxPrP mice. Fluorescence micrographs of Syn^Cre^;loxPrP and loxPrP COCS infected with prions or exposed to noninfectious brain homogenate (NBH). Slices were cultured for 56 days. Purkinje cells were identified by calbindin immunostaining. Calbindin and NeuN morphometry indicates degeneration of both granule cells and Purkinje cells in prion-infected Syn^Cre^;loxPrP mice. Each dot represents a cerebellar slice; n = 9. One-way ANOVA followed by Bonferroni’s post-hoc test was used for statistical analysis; ** p<0.01; *** p<0.001; ****p<0.0001. B: Prion mimetic anti PrP antibody POM1 triggers neuronal loss in COCS of PrP^C^ overexpressing mice (*tga20*). *tga20* COCS exposed to POM1 or IgG control and cultured for 10 days were stained for NeuN (green), calbindin (red) and DAPI (blue). NeuN morphometry revealed significant neurodegeneration in POM1 treated COCS. Each dot represents a single slice (n=9). One-way ANOVA followed by Bonferroni’s post-hoc test was used for statistical analysis; ** p<0.01; *** p<0.001; **** p<0.0001. C: Neurotoxic anti-PrP antibody POM1 does not trigger neuronal loss in COCS of GFAP^Cre^;loxPrP mice. Fluorescence micrographs of GFAP^Cre^;loxPrP+ and control GFAP^Cre^;loxPrP COCS exposed to POM1, or IgG control, and cultured for 10 days. COCS were stained for NeuN (green), calbindin (red) and DAPI (blue). Calbindin and NeuN morphometry indicated no neurodegeneration in POM1 treated GFAP^Cre^;loxPrP COCS. Each dot represents a cerebellar slice; n=9. One-way ANOVA followed by Bonferroni’s post-hoc test was used for statistical analysis; n.s.: not significant.

To further challenge the conclusion that astrocyte-restricted PrP^C^ expression does not restore prion-dependent neurodegeneration, we generated COCS from 9-day old pups of GFAP^Cre^;loxPrP and as controls from GFAP^Cre^ and *tg*a*20* mice (Fischer et al., 1996) which overexpress PrP^C^ almost ubiquitously. COCS were treated with the prion-mimetic antibody POM1 or with pooled mouse IgG, fixed after 10 days of treatment, and immunostained with NeuN and calbindin. COCS from *tg*a*20* mice, but not from GFAP^Cre^;loxPrP and GFAP^Cre^ mice, showed conspicuous neuronal loss (Figure 4B-C).

### Mice with neuron or astrocyte restricted PrP^C^ expression do not activate microglia upon prion infection

Prion diseases typically feature extreme activation and proliferation of microglia to an extent rarely seen in any other brain diseases. We assessed the status of microglia and astrocytes by immunohistochemistry for Iba1 and GFAP on hippocampal brain sections of terminally scrapie-sick wild-type mice and Syn^Cre^;loxPrP mice. For control we used loxPrP mice. While wild-type mice showed a high microglia density, we were surprised to find that Syn^Cre^;loxPrP mice did not show more microglial activation and astrogliosis than control mice (Figure 5A-C) despite being terminally scrapie-sick. We performed the same analysis in prion-infected GFAP^Cre^;loxPrP mice. Also here, the staining of cortical sections with anti-Iba1 and anti-GFAP did not reveal any microglia activation or astrogliosis beyond the baseline of control mice (GFAP^Cre-^;loxPrP, GFAP^Cre^;loxPrP^-^), whereas wild-type mice showed a brisk enhancement of Iba1 and GFAP immunoreactivity (Figure 5A-C).

**Figure 5:**
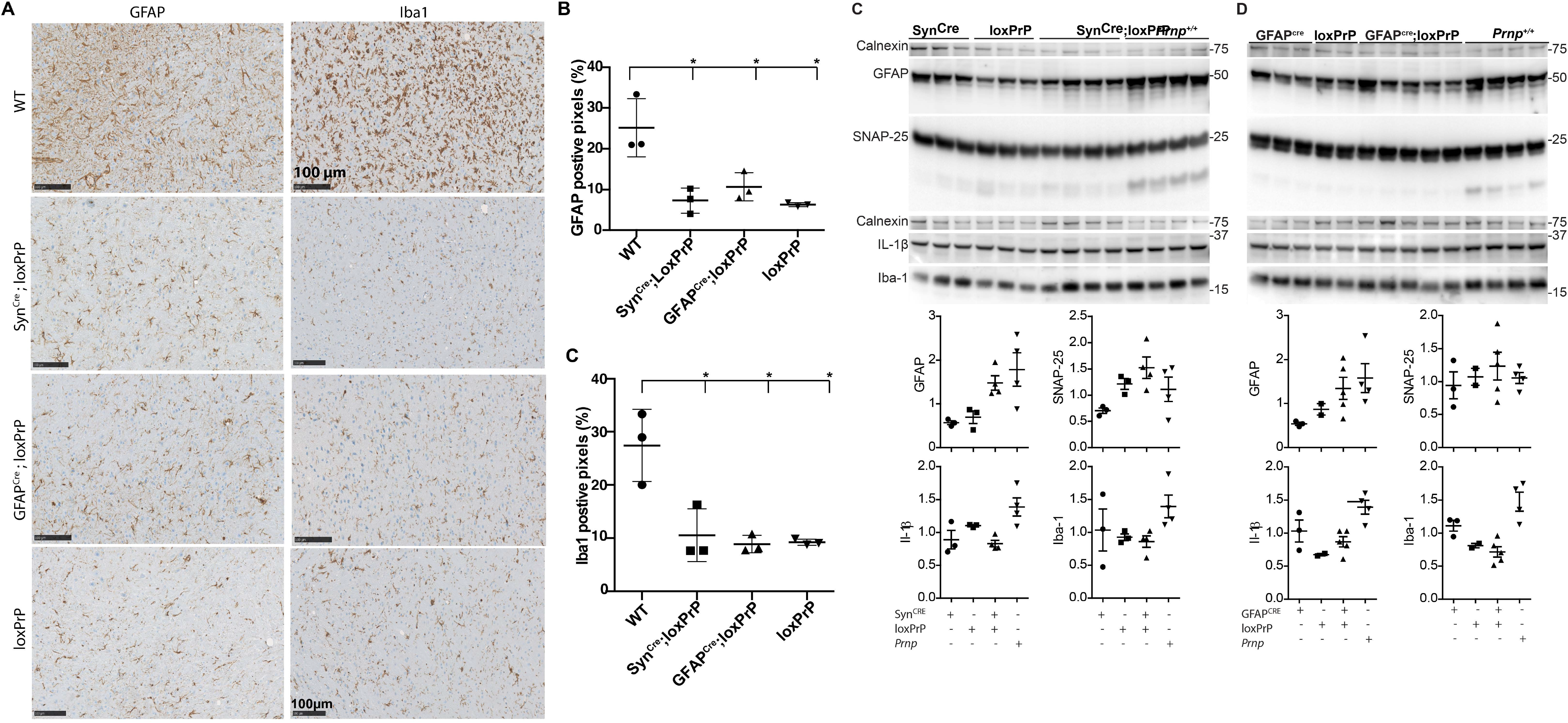
Prion infection of neuron and astrocyte-specific PrP expressors does not induce microglia activation. **A:** Microglia and astrocyte activation was analyzed by immunohistochemistry on hippocampal sections of prioninoculated Syn^Cre^;loxPrP (379 dpi), GFAP^Cre^;loxPrP (627 days), loxPrP (504 dpi) and *Prnp^+/+^* (184 dpi) mice. Sections were stained for Iba-1 and GFAP. **B-C:** Quantification of GFAP^+^ **(B)** and Iba1^+^ **(C)** pixels in a cross-section of complete mouse brain. Expression of both markers was conspicuously increased in *Prnp^+/+^* mice, whereas Syn^Cre^;loxPrP and GFAP^Cre^;loxPrP revealed significantly lower amounts of GFAP and Iba1. **C-D:** Western blot analysis and related quantification of GFAP, SNAP-25, IL1-Beta, and Iba-1 in prion inoculated Syn^Cre^;loxPrP mice (C) and GFAP^Cre^;loxPrP mice (D) versus controls. Calnexin was used for loading control. Only SNAP-25 showed elevated expression in prion infected Syn^Cre^;loxPrP mice whereas all other markers of microglial activation remained unaltered.

### Molecular changes associated with microglial activation remain unaltered in mice with cell type restricted PrP^C^ upon prion infection

Microglial changes typically precede the onset of the clinical signs of the disease, and are accompanied by the expression of pro-inflammatory cytokines such as TNFa, IL1α and IL1β. Hemispheric brain lysates from the same prion-infected mice as above were then subjected to Western blotting to monitor the expression of SNAP25 (a presynaptic protein engulfed by microglia previously shown to be reduced in prion infections before the onset of motor defects (Moreno et al., 2012)), GFAP, IL-1β and Iba-1. Terminally scrapie-sick wild-type mice showed a substantial increase in the expression of GFAP, Iba-1 and IL-1β, whereas Syn^Cre^;loxPrP and GFAP^Cre^;loxPrP lysates were similar to the negative controls. Only SNAP-25 was slightly elevated in Syn^Cre^;loxPrP mice (Figure 6A-B). These results indicate that microglial activation is not induced by prion infection in mice expressing PrP^C^ only in neurons or astrocytes.

**Figure 6:**
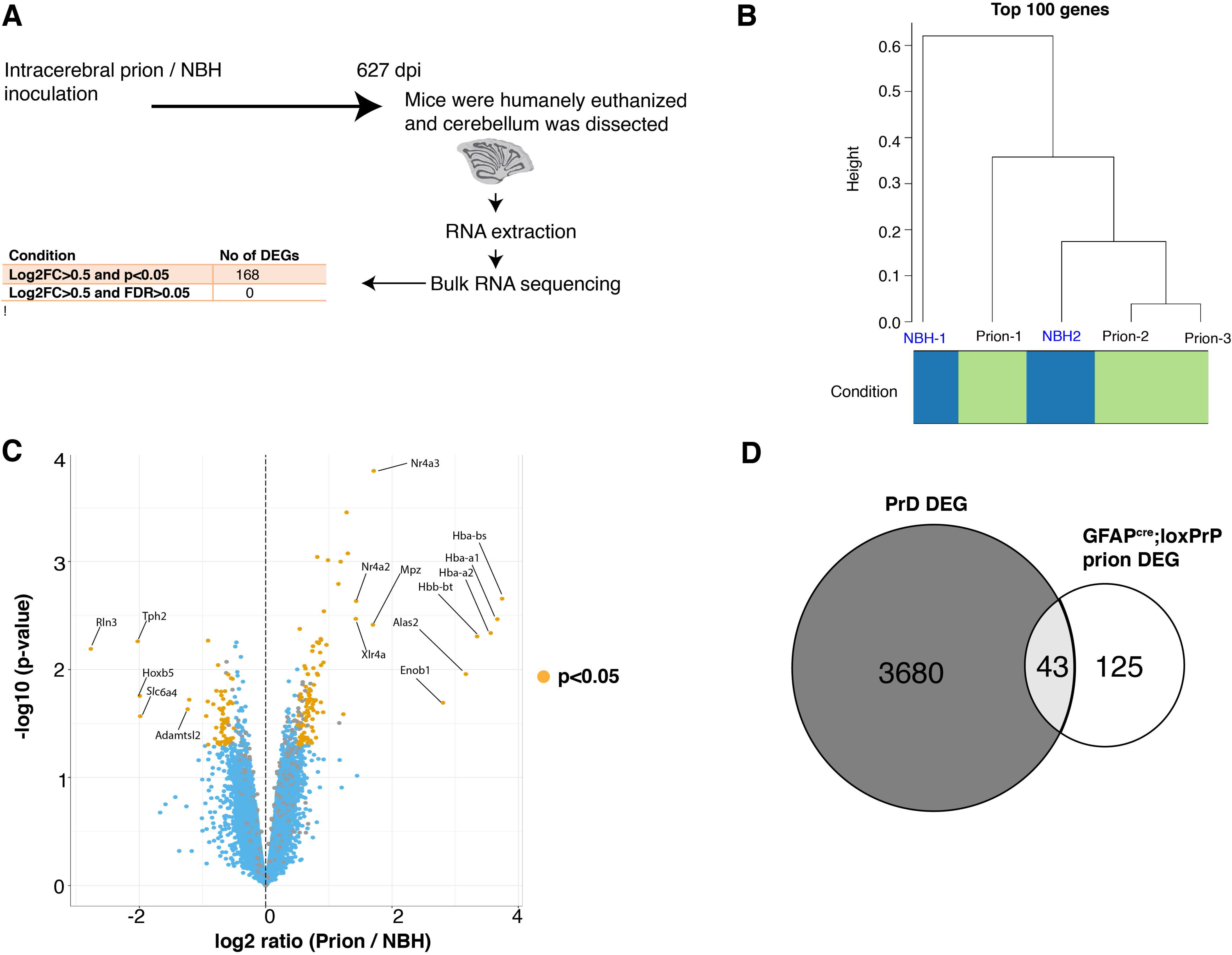
Prion disease specific transcriptional signature is absent in astrocyte specific PrP^C^ expressors infected with prions. **A:** Schematic representation of the sample collection and bulk RNA sequencing after prion or NBH inoculation of GFAP^Cre^;loxPrP mice. The number of differentially expressed genes (DEGs) and the filtering criteria used to derive them are indicated. **B:** Hierarchical clustering analysis based on the 100 genes with highest variance across all the samples did not reveal a separation between NBH and prion infected mice (n = 3 for prion inoculations and n = 2 for NBH injections). **C:** Volcano plot showing differentially expressed genes in the prion inoculated GFAP^Cre^;loxPrP mice compared to the NBH injected counterparts. Genes with Log_2_FC>0.5 and p-value<0.05 were considered as DEGs. The identities of the top 10 upregulated and top 5 downregulated genes are reported in the plot. Log2FC: log 2 fold change. **D:** Intersection between the DEGs observed during the progression of prion disease in C57BL6/J mice (3723 genes) and DEGs in prion inoculated GFAP^Cre^;loxPrP mice (168 genes). The intersection consists of only 43 genes, suggesting that prioninfected GFAP^Cre^;loxPrP mice do not exhibit any prion-specific transcriptomic signature.

### Prion infection does not elicit its transcriptional signature in mice expressing astrocyte-restricted PrP^C^

Previous studies have longitudinally mapped transcriptional changes associated with prion disease in mice, and revealed changes in gene expression from as early as 4 weeks post prion infection (Sorce et al., 2020). We asked if GFAP^Cre^;loxPrP mice, which do not develop clinical signs after prion infection, exhibit transcriptional changes upon prion infection. Three GFAP^Cre^;loxPrP mice were intracerebrally inoculated with RML6 prions; for control we inoculated two GFAP^Cre^;loxPrP mice with non-infectious brain homogenate (NBH). Prion-infected mice did not manifest prion disease and were humanely euthanized along with the NBH treated mice at approximately 627 dpi. RNA was isolated from cerebella of these mice and processed for RNA sequencing (Figure 6A). Unsupervised clustering of the 100 genes with highest variance across all the samples did not reveal any separation between the prion infected and NBH treated samples (Figure 6B). Data analysis to identify differentially expressed genes (DEGs) in prion inoculated vs NBH treated mice revealed absence of any DEGs when a filter of Log_2_FC>0.5 and FDR<0.05 was applied (Figure 6A). We then relaxed the stringency by applying a filter of Log_2_FC>0.5 and p-value<0.05. This revealed 168 DEGs in prion inoculated GFAP^Cre^;loxPrP mice. Of the 168 DEGs, 104 genes were upregulated, and 64 genes were downregulated (Figure 6C, Table S1). Functional gene ontology studies revealed upregulated genes were enriched in biological processes associated with hormonal response pathways (Figure S1C) whereas the downregulated genes were not enriched in any pathway. A total of 3723 genes were previously shown to have their expression, altered at least at one measured time point during the course of prion disease in C57BL6/J mice inoculated with RML6 prions. We next tested if any of the DEGs from the prion inoculated GFAP^Cre^;loxPrP mice overlapped with the DEGs observed during the progression of prion disease in C57BL6/J mice. Intersection plots revealed that 43 genes with altered expression in GFAPcre;loxPrP mice were also differentially expressed during the prion disease (Figure 6D, Table S2). These 43 DEGs however did not correlate with any particular time point during the progression of prion disease. We deduce that PrP^Sc^ produced by astrocytes does not induce the transcriptional changes typical of prion diseases.

## Discussion

While many transgenic mouse lines with tissue-specific PrP^C^ expression have been generated in the past two decades, we still lack a flexible generic model system allowing for expression in any given cell type under highly controlled conditions. What is worse, several of the transgenic models alleged to display cell-specific PrP^C^ expression in reality exhibit illegitimate expression in ectopic compartments: the widely used neuron specific enolase (NSE) promoter can be expressed in glial cells (Kugler et al., 2003) and earlier versions of a GFAP-driven construct are expressed in neurons (Marino et al., 2000; Zhuo et al., 2001). This situation has motivated us to generate the lines described in this study. We opted to use the CAG-CAT system (Araki et al., 1995) and in its native state, the CAG-CAT transgene leads to sustained and ubiquitous expression of chloramphenicol acetyl transferase (CAT), whose presence can be easily monitored with an enzyme activity assay. Upon CRE-mediated recombination, the loxP-flanked CAT minigene and its polyadenylation signal are excised, and transcription of PrP^C^ is enabled. We tested the system with two CRE-expressor crossings directing PrP^C^ expression to neurons and astrocytes. We chose the Synapsin-1 promoter as it is selectively, but broadly, in many types of neurons. Conversely, the mGFAP line 77.6 used in the current study has previously been shown to express Cre exclusively in astrocytes (Gregorian et al., 2009).

The data reported here enable new insights into two aspects of prion pathology: cell type-specific replication and cell type-specific toxicity. The expression of PrP^C^ is a necessary but insufficient prerequisite to prion replication, and multiple cell types replicate prions during their journey from the periphery to the brain. However, many tissues do not replicate prions despite expression of PrP^C^. While the replication competence of neurons is well established, the data on astrocytes have remained controversial. One of us (AA) reported that mice expressing hamster PrP^C^ off a GFAP promoter would replicate prions and develop disease (Raeber et al., 1997). However, a similar construct was found to be active ectopically in neurons (Marino et al., 2000), raising doubts on the specificity of the previous report. The specificity of transgenic mice described show is arguably more stringent. Although we were surprised to find that these mice did not develop any histological signs of neuroinflammation and clinical disease, we confirmed that astrocytes are capable of prion replication and PrP^Sc^ deposition in vivo.

The terminal stage of prion diseases is typically characterized by extensive neuronal loss. This observation has led to the implicit assumption that neurons are the primary targets, and possibly the primary driver, of prion diseases. An impressive panoply of evidence has accumulated that neuronal damage is crucially dependent on expression of PrP^C^ by the neurons targeted by prions. The first hint came from grafting PrP^C^-expressing neural tissue into PrP^C^-deficient mice. After prion infection, grafts developed all the pathological features of prion disease, yet no damage was observed in the adjoining tissue lacking PrP^C^ despite PrP^Sc^ deposition loss (Brandner et al., 2000; Glatzel and Aguzzi, 2000). An additional line of evidence came from partial depletion of neuronal PrP^C^, which decelerated the development of disease despite deposition of PrP^Sc^ (Mallucci et al., 2003). Furthermore, transgenic mice expressing a hamster prion protein in neurons became susceptible to prion disease after exposure to hamster prion, suggesting that mere expression of PrP^C^ in neurons suffices to develop prion disease (Race et al., 1995).

On the other hand, a growing number of observations is challenging the perception that neurons are the sole, or even the dominant, cellular actor in prion pathology. Recently concluded longitudinal transcriptomic study mapping the changes in gene expression during the course of prion infection has revealed that glial perturbations occur simultaneously to the onset of clinical disease. Surprisingly, no changes in the expression of neuron-enriched transcript were detectable at this time point. Instead, suppression of neuronal transcripts was only observed at the terminal stage of disease (Sorce et al., 2020). The onset of changes associated with actively translating genes during the course of prion infection also revealed glial perturbation as the major contributors in driving the course of prion disease (Scheckel et al., 2020). More importantly, disease associated microglia (DAM) genes and A1 astrocytes, both of which are renowned glia signatures in several neurodegenerative diseases were upregulated in prion disease. Overall, these studies revealed an important role for non-neuronal cell types in the manifestation of prion disease.

### Contribution of cell type specific PrP^C^ to prion disease

Prion inoculated Syn^Cre^;loxPrP mice displayed all the characteristic histopathological and clinical features of prion disease. The incubation time was much longer than that of wild-type mice (median survival: 435 vs 196 days), which may be explained by the lower total brain PrP^C^ concentration.

Although the mechanisms responsible for neuronal death are yet to be elucidated it is evident that PrP^Sc^ deposits from non-neuronal counterparts could accelerate the conversion of neuronal PrP^C^ and thereby aggravating neuronal loss. Interestingly, these mice showed no activation of microglia and astrocytes and this further confirms the hypothesis that glial activation drives the progression of the disease and in its absence the onset of the disease follows a much longer time course. The Syn^Cre^;loxPrP^+^ mice line delinks glial activation from neuronal death an could be potentially used as a model system to study the mechanisms associated with neuronal death in prion disease.

Among non-neuronal cells, astrocytes are the cell type producing highest amount of PrP^C^ (Hartmann et al., 2013). Previous studies have shown astrocytes not only capable of replicating and propagating prions but also can deposit PrP^Sc^ aggregates (Raeber et al., 1997). Transgenic mice expressing PrP^C^ exclusively in astrocytes (GFAP^Cre;^loxPrP) do not develop prion disease and do not show any of the classical pathological features apart from deposition of PrP^Sc^. These mice also fail to recapitulate glia activation or upregulate any of the inflammation markers. RNA sequencing data corroborated the absence of molecular markers of disease. Differentially expressed genes became identifiable only when the stringency of the analysis was substantially relaxed, and even then they did not appear to be enriched in any specific cell types or pathways. We conclude that astrocyte-selective prion infection has remarkably bland effects on the brain.

Previously, mice expressing PrP^C^ under the expression of GFAP promoter were shown to replicate prions but did not develop the disease (Diedrich et al., 1991). The capability of astrocytes to replicate prions was however questioned because the GFAP promoter fragment used to generate the mice also exhibited partial neuronal expression (Marino et al., 2000; Zhuo et al., 2001). Using the newly generated mouse model, we have now validated that astrocytes are indeed capable of replicating prions and further vindicates a recent iPSC derived astrocyte cell culture model system which successfully replicated CJD prions (Krejciova et al., 2017). Recent studies have reported transport of PrP^Sc^ deposits from astrocytes to neurons predicting the possibility that astrocytic PrP^Sc^ is toxic to neurons (Victoria et al., 2016). Our current mouse model system proves that this is indeed not the case and in spite of astrocytic PrP^Sc^ accumulation there is apparent neurotoxicity in the mice.

In the future, these newly generated mice lines can help answering some of the long-standing questions in the prion field and may represent ideal tools to delineate the role and contribution of each of the cell types in manifestation of prion disease.

## Material and Methods

### Mice

Animal welfare and experimental procedure on the mice were performed according to the “Swiss Ethical Principles and Guidelines for Experiments on Animals” and approved by the Veterinary office of the Canton of Zurich (permit 90/2013). All efforts were made to minimize the suffering and reduce the number of animals used for the experiments. Mice were bred and housed in special hygienic grade facilities and housed in small groups (max 5 per cage) under a 12h light / 12h dark cycle with sterilized food (Kilba No.3431, Provimi Kilba Kaiseraugst, Switzerland) and water *ad libitum*. Prion inoculated mice were regularly monitored according to the standard operating procedures approved by the Veterinary office and mice were humanely sacrificed once the termination criteria were reached.

#### Generation of CAG-CAT-PrP transgenic mice

For the generation of mice carrying a *loxP*-flanked stop cassette followed by the mouse Prnp coding sequence under the CAG promoter, we used the pCAG– *lox*P–CAT–*lox*P vector (kind gift of Dr. Kimi Araki, Kumamoto University, Japan), where CAG is the CMV immediate early enhancer-chicken β-actin hybrid promoter and CAT the chloramphenicol acetyl transferase gene (Araki et al., 1995). Cloning by T4 DNA ligase (New England Biolaboratories, Ipswich, MA, United States) was performed after *Eco*RV (New England Biolaboratories, Ipswich, MA, United States) digestion of both the PCR-amplified *Prnp* cDNA and the pCAG–*lox*P–CAT–*lox*P vector. The *Kpn*I-*Sac*I (New England Biolaboratories, Ipswich, MA, United States) linearized CAG-CAT-Prnp transgene was purified and microinjected at the transgenic facility of the University Hospital Zurich into one cell-stage fertilized embryos from Prnp^ZH1/ZH1^ mice. Transgenic founders were identified by PCR using the following primers: Forward: AAC GCC AAT AGG GAC TTT CC; Reverse: ATG GGG AGA GTG AAG CAG AA (actin primers used as internal control for PCR: fwd -TGT TAC CAA CTG GGA CGA CA; rev- GAC ATG CAA GGA GTG CAA GA). Following assessment of germline transmission and establishment of the colonies, highest expresser line (Line 211) was selected based on the levels of CAT expression which were determined in brain tissue homogenates of two-month old CAG-CAT-Prnp mice using the CAT ELISA kit (Roche, Basel, Switzerland) according to manufacturer’s instructions. Selected line was backcrossed into C57BL6/J background and Prnp^ZH3/ZH3^ for 9 generations.

To generate mice expressing Cre recombinase under the control of cell type specific promoters in ZH3 background: Synapsin1 Cre (B6.Cg-Tg(Syn1-Cre)671Jxm/J(#003966), Plp1 CreER (B6.Cg-Tg(Plp1-Cre/ERT)3Pop/J(#005975) and GFAP Cre mice (B6.Cg-Tg(Gfap-Cre)77.6Mvs/2J(#024098) were bred with ZH3. Cell type specific Cre expressors in ZH3 background were then then crossed with Line 211 to generate mice lines expressing PrP^C^ exclusively in subset of neurons (Syn^Cre^;loxPrP), astrocytes (GFAP^Cre^;loxPrP) and oligodendrocytes/myelinating Schwann cells (Plp^Cre^;loxPrP). Other mice lines used in the current study are: C57BL6/6J and tga20 mice (B6.Cg-Tg(Prnp)a20Cwe).

#### Prion inoculations

RML6 brain homogenates were prepared from the brains of terminally sick CD1 mice infected with RML6 prions. Brain homogenates were prepared in PBS (+5%BSA). For control inoculations brain homogenates from healthy CD1 mice were used and they are refereed to as non-infectious brain homogenate (NBH). 30μl of RML6 (dose corresponding to 3×10^5^ LD_50_) or NBH lysate was injected intracerebrally into 6-8 week old mice. Scrapie was diagnosed according to clinical criteria (ataxia, kyphosis, priapism, and hind leg paresis). Mice were sacrificed on the day of onset of terminal clinical signs of scrapie and NBH inoculated mice were sacrificed approximately at the same time. In case of mice that did not manifest prion disease, they were sacrificed approximately after 550-630 days post inoculations.

### RNA Sequencing and Data Analysis

RNA extraction from cerebella of mice was performed as described previously (Sorce et al., 2020). Data analysis and visualizations were performed using Sushi data analysis framework provided by Functional genomics center Zurich (FGCZ) from University of Zurich. Quality control and data analysis was performed as described previously (Henzi et al., 2020). Differential gene expression was performed using Edge R (version 3.0) (Robinson et al., 2010) and any gene with Log2FC>0.5 and p-value<0.05 was considered to be differentially expressed. Data intersection with genes dysregulated during the course of prion infection was performed using Multiple list comparator from www.molbiotools.com.

### Cerebellar Organotypic Cultured Slices (COCS)

350μm thick COCS were prepared from 9-12 day old mice pups as described previously (Falsig and Aguzzi, 2008). COCS cultures were maintained in a standard incubator (37°C, 5% CO2, 95% humidity) and the medium was replenished three times per week.

#### Prion inoculations of COCS

Freshly prepared COCS were inoculated with Rocky mountain laboratory strain 6 prions (RML6) or as a control with either noninfectious brain homogenate (NBH) as described previously. Slices were maintained f for a further 45 days followed by fixation with 4% paraformaldehyde and staining with NeuN (to label cerebellar granule neurons), calbindin (to label Purkinjee cells) and DAPI (to label nuclei). NeuN and calbindin morphometry was analyzed using analySISvc5.0 software and neurotoxicity was defined as significant loss of NeuN or calbindin positive neuronal layer loss over NBH treatment.

#### POM1 treatment of COCS

Toxicity in COCS was induced by treatment with anti-PrP^C^ antibody (POM1) targeting the globular domain of the protein as described previously (Sonati et al., 2013). Anti-globular domain ligands. COCS were treated with wither POM1 (67nM) or as control with IgG for 10 days after a 14-day recovery period of initial gliosis due to tissue preparation. COCS were fixed, imaged and analyzed as described for the prion inoculated slices. Antibody treatment was randomly assigned to individual wells.

### Western Blots

Mice brains (RML6 and NBH inoculated) were homogenized using Ribolyser for 5 min in 10 vol of lysis buffer (0.5% Nonidet P-40, 0.5% 3-[(3-cholamidopropyl)dimethylammonio]-1-propanesulfonate (CHAPS), protease inhibitors (complete Mini, Roche), phosphatase inhibitors (PhosphoSTOP, Roche) in PBS, and centrifuged at 1000g for 5 min at 4°C to remove debris prior to analysis by SDS-PAGE (Novex NuPAGE 10% Bis-Tris Gels). After electrophoresis, gel was transferred to iBlot I (Invitrogen) and transferred onto nitrocellulose membranes. Membranes were blocked in 5% Sureblock for 1 h at room temperature followed by incubation with primary antibody overnight. Membrane was washed 3× (15 min each) with PBS-Triton (0.2%) followed by incubation with HRP-tagged secondary antibody (Peroxidase-Goat Anti-Mouse IgG (H+L) (#62-6520) or Peroxidase-Goat Anti-Rabbit IgG (H+L) (#111.035.045); 1h at room temperature) and further washes (3×, 10 min). Membrane was developed with Luminata Crescendo (Millipore) and images were acquired using LAS-3000 Imaging system from FUJI. Densitometry analysis was performed using Quantity One software (BioRAD) and data was plotted using Graphpad software.

### Immunohistochemistry

Brain tissues were fixed in formalin and treated with concentrated formic acid to inactivate prions. 2μm thick sections were prepared from these brains and deparaffinized using graded alcohols and then subjected to antigen retrieval using 10mM citrate buffer (pH 6). Astrogliosis, Microgliosis and the presence of protease resistant-prion deposits were visualized by staining brain sections with GFAP (1∶1000, Agilent technologies), IBA1 (1∶2500, WAKO) and the SAF84 antibody (1∶200, SPI bio) respectively on a NexES immunohistochemistry robot (Ventana instruments) using an IVIEW DAB Detection Kit (Ventana). Sections were also counterstained with Hematoxylin and eosin when appropriate. Images were acquired using NonoZoomer scanner (Hamamatsu Photonics) and visualized using NanoZoomer digital pathology software (NDPview; Hamamatsu Photonics). Images were acquired using Olympus BX61 Upright fluorescent microscope. Quantifications of IBA1, GFAP staining was performed on entire brain section from the acquired images using Image J.

### Antibodies

The following antibodies are used in the current study.

| Antibody | Source | Cat.No |
| --- | --- | --- |
| anti-SNAP25 antibody | Abcam | ab5666 |
| anti-GFAP | Agilent technologies | Z03344 |
| anti-Iba1 | WAKO | 019-19741 |
| anti-SAF84 | Bertin bioreagent | A03208 |
| anti-NeuN | Millipore | MAB377X |
| anti-calbindin | Abcam | ab108404 |
| anti-calnexin | Enzo Life sciences | ADI-SPA-865-D |
| anti IL1β | Abcam | ab9722 |
| anti-POM1 antibody | Aguzzi Lab | (Polymenidou et al.,<br>2008) |
| anti-actin | Millipore | MAB1501R |

### Statistical Analysis

Details of the type of statistical tests performed are described in the figure legends. Results are represented as mean of replicates (±SEM) Data was analyzed using GraphPad software.

## Acknowledgements

We thank Mirzet Delic and Paulina Pawlak for animal husbandry, Merve Avar for help with data analysis of RNA sequencing data. AA is the recipient of an Advanced Grant of the European Research Council, the Swiss National Foundation, the Nomis Foundation and SystemsX.ch. AS, SS and AKKL are recipients of a grant from the Synapsis Foundation.

## Author contributions

AA initiated and supervised the project, and wrote the manuscript. AKKL performed RNA sequencing experiment, analyzed all the data and wrote the manuscript. AS performed slice cultures, Immunohistochemistry, ELISA, western blots and analyzed the data. SS contributed to generation of mice lines, immunohistochemistry, ELISA and analyzed the data. MN contributed to generation of mice lines. OR, JG, PS, RM provided the technical help in performing all the experiments in the manuscript. PP performed microinjections and contributed to the generation of mice lines. All authors have read and approved the final version of the manuscript.

## Supplemental Figures

**Figure S1: A:** Polymerase chain reaction (PCR) performed on the tail biopsies of seven transgenic founder mice revealed successful integration of CAT-PrP transgene. For control, we used cDNA of CAT-PrP and DNA extracted from C57BL6/J mouse tail biopsies. Actin was also simultaneously amplified to ensure that PCR mix worked efficiently. Samples were migrated on 2% Agarose in Tris-EDTA buffer for electrophoresis. Lines 208, 214 and 215 failed to transmit the transgene to F1 generation. **B:** Immunohistochemistry on the hippocampal sections of Syn^Cre^;loxPrP (337 dpi), GFAP^Cre^;loxPrP (574 days) inoculated with NBH, to monitor the activation of microglia (Iba1) and astrocytes (GFAP). Staining revealed no activation of microglia and astrocytes in both sections. **C:** Biological process enrichment obtained by gene ontology analysis on the upregulated genes in prion infected GFAP^Cre^;loxPrP mice revealed that most upregulated genes were involved pathways associated with hormone response.

**Figure S2:** Uncropped Western blots

All Western blots pertaining to this study are shown here in their original format as outputted by the blot imager.

**Table S1:** Table including detailed RNA Sequencing results including the differentially expressed genes, p-values, FDR values in prion infected GFAP^Cre^;loxPrP mice.

**Table S2:** Table showing differentially expressed genes that are common to prion disease (PD) and prion infected GFAP^Cre^;loxPrP mice (GFAP^Cre^;loxPrP prion).

