## Supplementary Figure 2 for "Glial activation in prion diseases is strictly nonautonomous and requires neuronal PrP^Sc^"

### Figure 3B

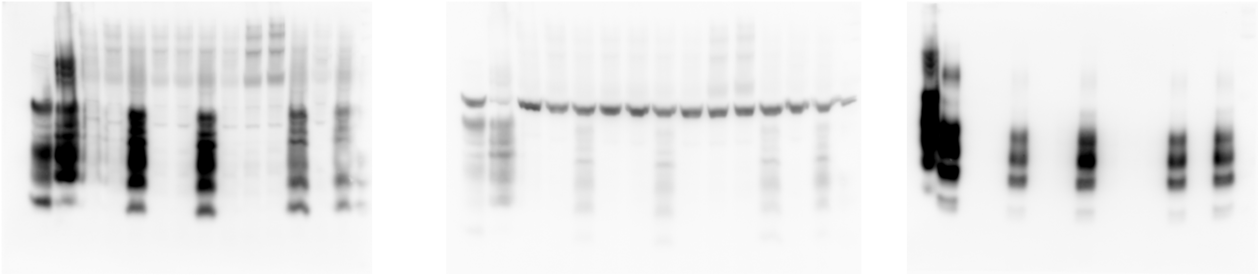

Undigested PrP | Actin | PK digested PrP

### Figure 3D

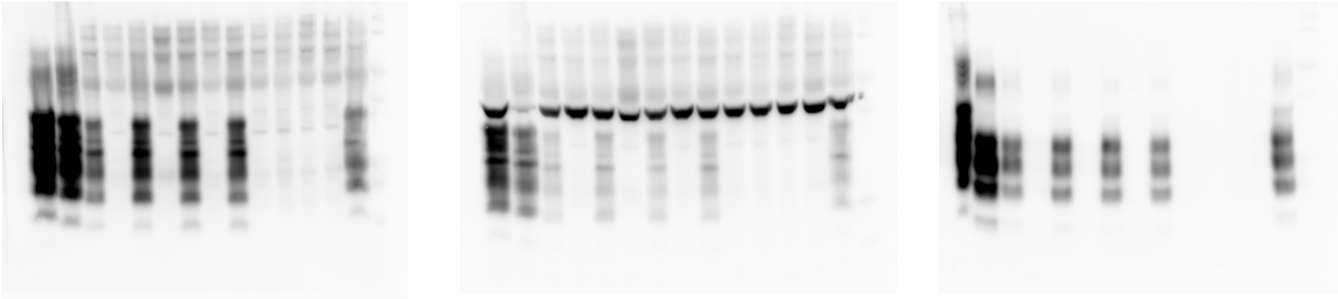

Undigested PrP | Actin | PK digested PrP

### Figure 5C

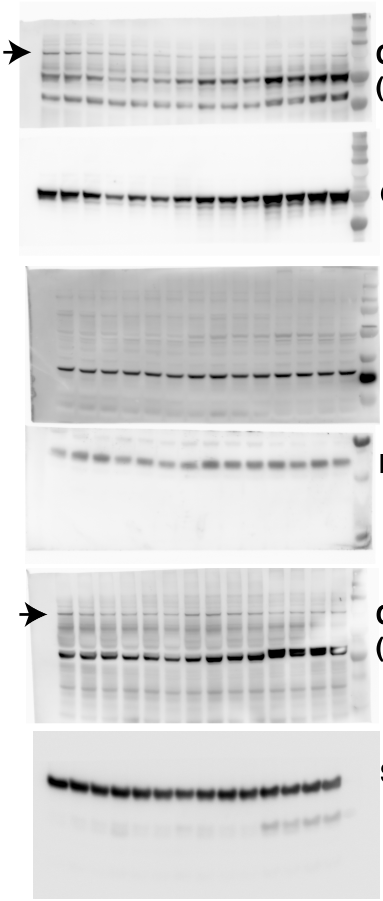


### Figure 5D

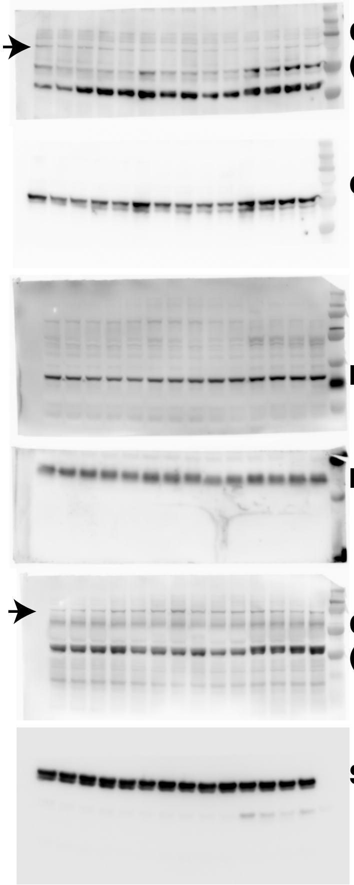
