## Supplementary figures and images for "Glial activation in prion diseases is strictly nonautonomous and requires neuronal PrP^Sc^"

### Supplementary Figure 1

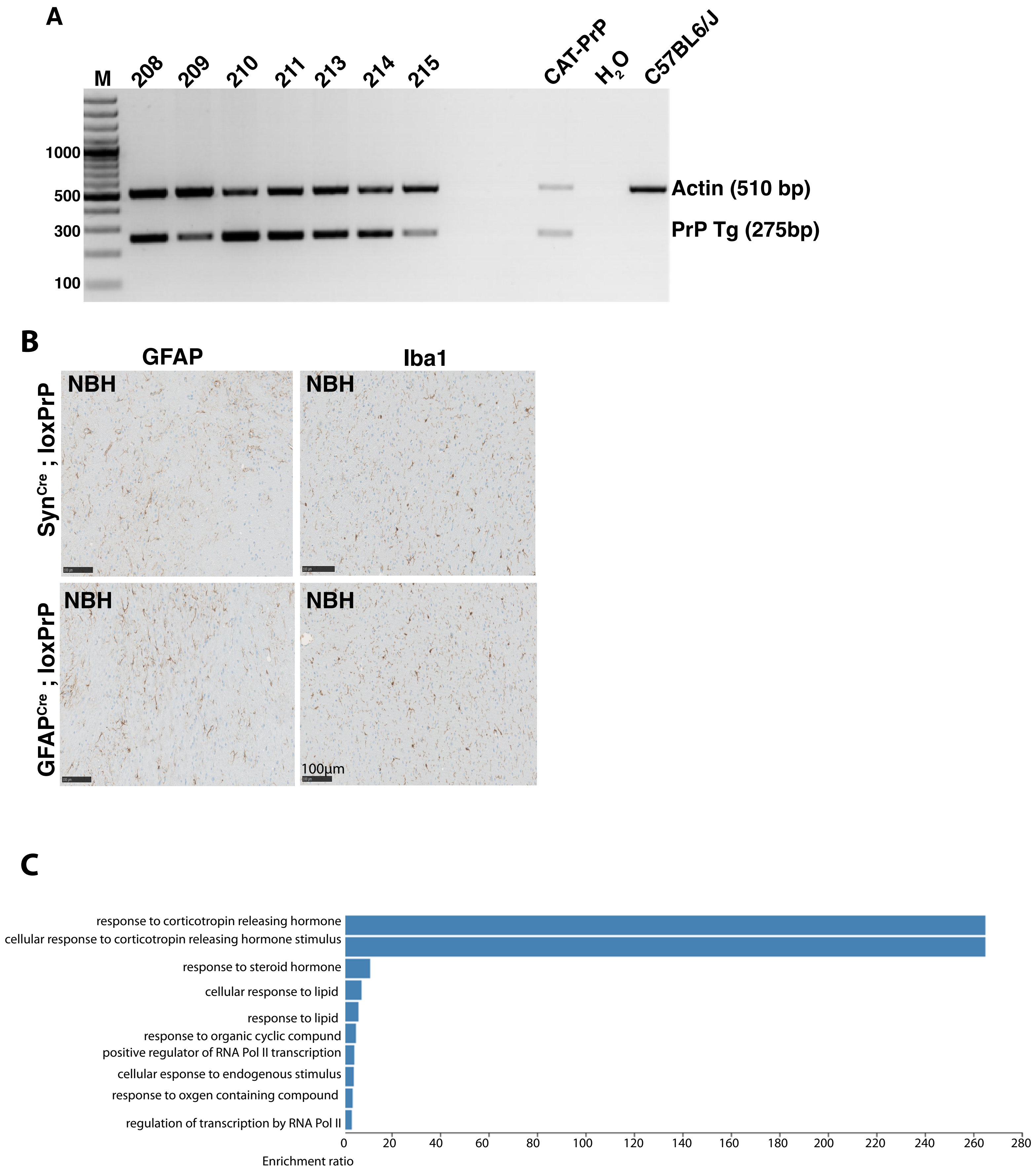


Senatore et al; Figure S1
